# SARS-CoV-2 fears green: the chlorophyll catabolite Pheophorbide a is a potent antiviral

**DOI:** 10.1101/2021.07.31.454592

**Authors:** Guillermo H. Jimenez-Aleman, Victoria Castro, Addis Longdaitsbehere, Marta Gutierrez-Rodríguez, Urtzi Garaigorta, Roberto Solano, Pablo Gastaminza

## Abstract

SARS-CoV-2 pandemic is having devastating consequences worldwide. Although vaccination advances at good pace, effectiveness against emerging variants is unpredictable. The virus has displayed a remarkable resistance to treatments and no drugs have been proved fully effective against Covid-19. Thus, despite the international efforts, there is still an urgent need for new potent and safe antivirals against SARS-CoV-2. Here we exploited the enormous potential of plant metabolism using the bryophyte *Marchantia polymorpha* and identified a potent SARS-CoV-2 antiviral, following a bioactivity-guided fractionation and mass-spectrometry approach. We found that the chlorophyll derivative Pheophorbide a (PheoA), a porphyrin compound similar to animal Protoporphyrin IX, has an extraordinary antiviral activity against SARS-CoV-2 preventing infection of cultured monkey and human cells, without noticeable cytotoxicity. We also show that PheoA prevents coronavirus entry into the cells by directly targeting the viral particle. Besides SARS-CoV-2, PheoA also displayed a broad-spectrum antiviral activity against (+) strand RNA viral pathogens such as HCV, West Nile, and other coronaviruses, but not against (−) strand RNA viruses, such as VSV. Our results indicate that PheoA displays a remarkable potency and a satisfactory therapeutic index, which together with its previous use in photoactivable cancer therapy in humans, suggest that it may be considered as a potential candidate for antiviral therapy against SARS-CoV-2.

## Introduction

The pandemic caused by the severe acute respiratory syndrome coronavirus 2 (SARS-CoV-2) is having devastating consequences, with more than 196M infected people and over 4M deaths worldwide (July 2021; https://covid19.who.int/). Besides the humanitarian cost, this pandemic carries a tremendous negative economic impact, a huge challenge for any government to overcome. The coronavirus disease 2019 (Covid-19), the respiratory illness caused by SARS-CoV-2 (Genus betacoronavirus; Subgenus sarbecovirus), has displayed a remarkable resistance to treatments and no drugs have been proved fully effective against the virus. Moreover, the Covid-19 pandemic has made evident the need for a global strategy to fight similar situations that may appear in the future.

Current efforts to eradicate Covid-19 are focused on the development of vaccines and the search for antiviral lead compounds, mainly repurposing of existing drugs. Although the vaccination campaign seems to advance at good pace, its effectiveness against some of the present and future strains of SARS-CoV-2 is hard to predict due to the existence of different strains that could drastically reduce the vaccine efficiency^1^. In addition, the best anti-Covid-19 drugs approved so far (e.g. Remdesivir, Favipiravir or its derivative Avifavir, etc), have shown only a mild effect against the virus, slightly reducing hospitalization time of patients^2^. Other treatments, such as Chloroquine and Hydroxychloroquine appear to help at least a subgroup of patients, but, possible negative side effects of these drugs remain under investigation^3^. Although, some compounds (e.g., Aplidin, Mefloquine, Nelfinavir, Protoporphyrin IX and Verteporfin) have shown potential on *in vitro* assays, and some of them also in animal models^4–6^, there is an urgent need for new potent and safe antivirals against SARS-CoV-2. Noteworthy, new pathogens, including viruses, are expected to emerge in coming decades, which puts an enormous pressure on society in order to be ready to fight back future pandemics with the proper chemical, biological, and engineering tools, including effective new antivirals.

For centuries, medical needs of society have been widely covered by plants, which have an extremely rich metabolism that provides them with a wide repertoire of chemical weapons to cope with environmental biotic stresses, including viruses^7,8^. Originally recognized by traditional medicine, plants are the main source of compounds used today in pharmacology, from Aspirin (acetyl salicilate; from Salix sp.) to current anticancer drugs (e.g. Vinblastine and Vincristine from Vinca sp., or Taxol and Paclitaxel from Taxus baccata), simply to cite a few successful examples^9,10^. Therefore, the identification of new plant sources of enzymatic variants and metabolites is essential to the discovery of new drugs and their optimization by metabolic engineering. Aromatic and exotic vascular plants are commonly studied in order to identify pharmacologically interesting compounds. In contrast, the metabolic richness of bryophytes (non-vascular plants including mosses, liverworts and hornworts) has been little explored. Bryophytes are rarely attacked by pathogens (fungi, bacteria, viruses) or herbivores (insects, snails, mammals) in their natural habitats, which indicates that they are well protected by a potent arsenal of secondary defense metabolites. However, studies on their chemical constituents have been neglected until recently^11^. Indeed, only around 5% of bryophyte species have been metabolically explored, and results have shown an enormously rich diversity of secondary metabolites, particularly in liverworts^12^. Strikingly, more than 1600 terpenoids have been reported in liverworts, whereas only about 100 terpenoids have been identified in the medicinal plant *Cannabis sativa*^11,13–16^. More importantly, several liverwort species of the order Marchantiales, including *Marchantia polymorpha*, produce terpenoids and bisbibenzyls with enormous potential for pharmaceutical applications since they show remarkable antimicrobial, antioxidant, cytotoxic, anticancer and antiviral (anti-HIV) activities^11,13,15–18^. Therefore, we made use of our vast experience in vorology and Marchantia’s hormonal signaling and secondary metabolism in order to explore this plant’s potential as a source of antiviral metabolites, particularly, against the SARS-CoV-2 virus.

In this study, we employed a set of *Marchantia* wild type plants, and signalling and metabolic mutants to systematically study the pharmacological potential of this liverwort. We found that total extracts from all the plants displayed a remarkable antiviral activity against SARS-CoV-2. Using a bioactivity-guided chromatographic approach, in addition to mass-spectrometry (MS), we identified the antiviral metabolite as Pheophorbide a (PheoA), a porphyrin chlorophyll derivative very similar to animal Protoporphyrin IX, also described as an strong antiviral^19^. In contrast to Protoporphyrin IX, which produces prophyria in humans, PheoA, however, was non toxic at all tested concentration. We also found that PheoA has a broad-range antiviral activity against positive strand RNA (+RNA) viruses, and acts as a virucidal by directly acting on the viral particle. PheoA is additive to remdesivir, which, together with its low toxicity, suggest its potential as candidate for antiviral therapy against SARS-CoV-2.

## Results

### Crude extracts of M. polymorpha show anti-SARS-CoV-2 activity

In order to explore for the presence of anti-SARS-CoV-2 metabolites in *M. polymorpha*, we prepared crude extracts of two different *M. polymorpha* subspecies (subsp. *ruderalis* from Japan and subsp. *polymorpha*, from Spain). The anti-SARS-CoV-2 activity of the resulting crude extracts was tested in Vero E6 cell monolayers infected with the SARS-CoV-2 NL2020 strain (Figure 1A). Infection was carried out at a multiplicity of infection (MOI) of 0.001 for 72 h. In the absence of antiviral activity, SARS-CoV-2 infection triggers cell death of Vero E6 cells and results in loss of cell biomass in the well, which is readily visualized as a strong reduction in crystal violet staining (DMSO, Figure 1A). Treatment of the cells during infection with serial dilutions of remdesivir (used as positive control), the only clinically approved antiviral for treatment of Covid-19 patients, protected the cell monolayers down to its reported EC_50_ of 1.5 μM (Supplementary Figure S1). Similarly, treatment of cell cultures with Marchantia crude extracts resulted in cell protection against virus-induced cytopathic effect without any signs of cytotoxicity in a broad dilution range (Figure 1A, Ex1 and Ex2), suggesting the presence of one or more *Marchantia* metabolites with strong antiviral activity. Given that antiviral activity had been determined by an indirect measurement (Figure 1B), we set out to define if *Marchantia* extracts were indeed capable of interfering with viral spread in cell culture. Thus, Vero E6 cells were inoculated at MOI of 0.001 in the presence of control-solvent, remdesivir (25 μM) and a 1:800 (v/v) dilution of the *Marchantia* extracts. Viral RNA load, which in this experimental setup represents the degree of virus propagation, was determined 72 h post infection by RT-qPCR (Figure 1C). In the control, viral RNA accumulated six orders of magnitude above the assay background levels, whereas the viral RNA was undetectable in samples treated with the antiviral remdesivir. Importantly, in samples treated with *Marchantia* extracts, the viral RNA levels were comparable to those observed upon remdesivir treatment (Figure 1C). This outcome confirms the protective activity of the extracts observed in Figure 1A,B and suggest the presence of at least one antiviral compound.

**Figure 1:**
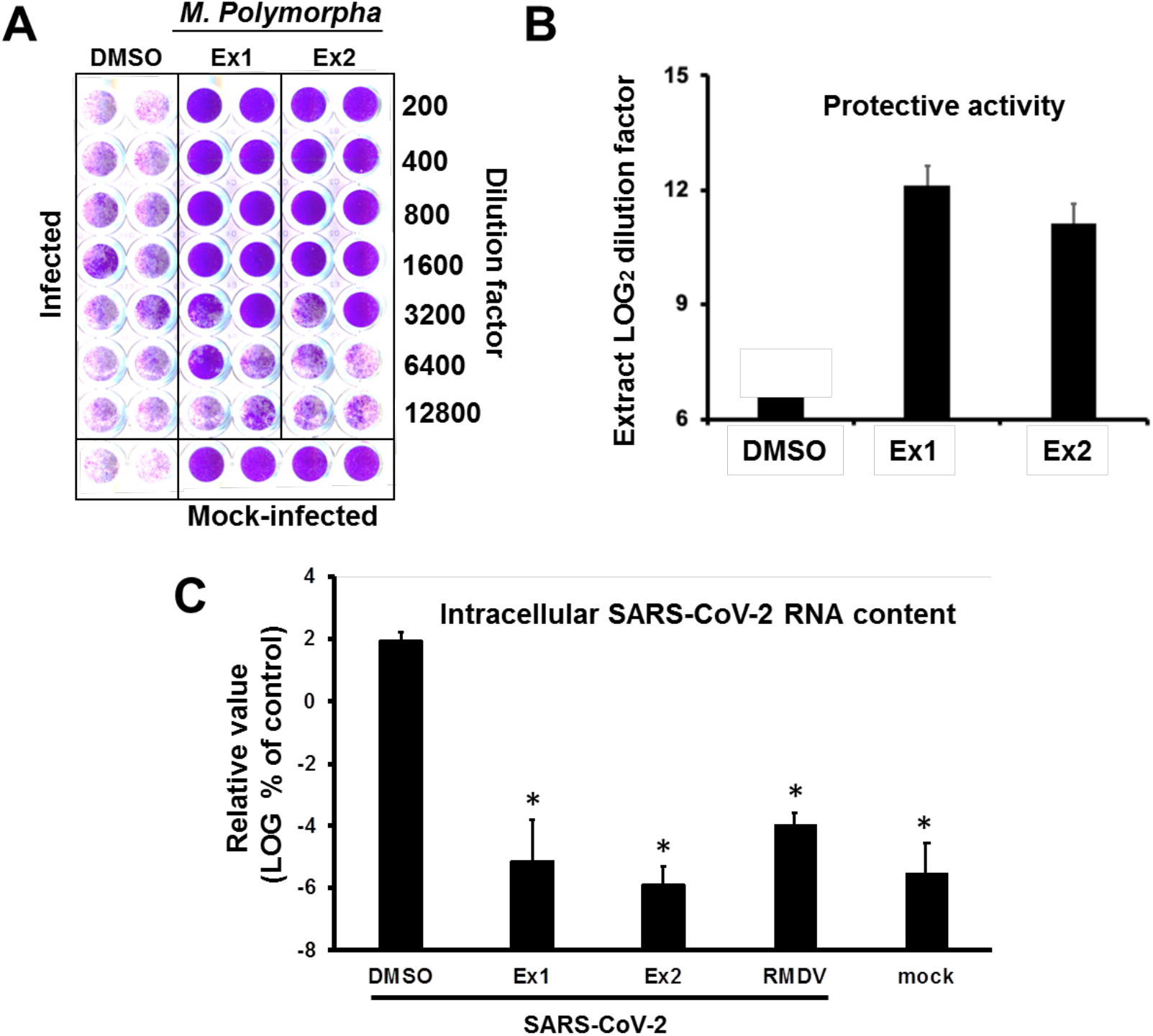
Marchantia polymorpha extracts interfere with SARS-CoV-2-induced cytopathic effect and virus propagation. (A-B) Vero E6 cells were inoculated with SARS-CoV-2 (MOI = 0.001) in the presence of serial 2-fold dilutions of crude extracts from two different *M. polymorpha* ecotypes, *ruderalis* (Ex1) and *polymorpha* (Ex2) and incubated for 72 h, time after which they were fixed and stained with a crystal violet solution. Mock-infected cells were used as the control of the integrity of the cell monolayer. A) Image of an experimental plate showing dose-dependent protection of *M. polymorpha* extracts in comparison with vehicle (DMSO)-treated cells. B) Numeric expression of the protective capacity as the log2 value of the highest dilution factor capable of full-monolayer protection. Data are shown as average and mean error of two biological replicates. (C) Vero E6 cells were inoculated with SARS-CoV-2 (MOI = 0.001) in the presence of vehicle (DMSO), remdesivir (RMDV; 25 mM) or a 1:800 (v/v) dilution of crude extracts. Parallel samples remained uninfected as control (mock). Total RNA was extracted 72 h post-infection and subjected to RT-qPCR to determine viral load. Normalized viral RNA levels are shown as percentage of the viral RNA found in vehicle-treated cells. Data are shown as mean (± SD) of three biological replicates. Statistical significance was estimated using one-way ANOVA and a Dunnet′s post-hoc test (*p<0.05).

### Extract bioactivity does not depend on plant’s secondary metabolism and is common to several plant species

Next, we explored whether the putative antiviral metabolite could belong to plant’s secondary metabolism. Jasmonates (JAs) are a family of oxylipin-derived phytohormones regulating many aspects of plant development and growth; as well as mediating defense responses through transcriptional activation of the secondary metabolism, which includes several classes of compounds such as alkaloids, terpenoids and flavonoids^20–22^. In *Marchantia*, secondary metabolites accumulate in specific organelles named oil bodies (OB), which are confined to scattered idioblastic OB cells distributed throughout the thallus^23^. Therefore, we tested extracts from *M. polymorpha* WT, Mp*coi1-2* [impaired in dn-OPDA perception, the active jasmonate in Marchantia^22^, thus, in defense metabolite induction], and Mp*c1hdz* plants (impaired in OB formation; MpC1HDZ is a transcription factor required for OB cells differentiation). Mp*c1hdz* mutants render plants defective in secondary metabolites, thus, susceptible to herbivory and microbes^24^. To our surprise, all Marchantia extracts, WT or mutant, showed similar antiviral activity (Figure 2), indicating that the active antiviral should not belong to the plants’s secondary metabolism. Indeed, data in Figure 2 suggests that the activity of extracts is due to the presence of a metabolite (or metabolites) that is constitutively synthesized and/or derived from the plant’s primary metabolism. Remarkably, regulation of primary metabolism is achieved in all plants by very similar conserved metabolic pathways^25,26^. Therefore, we tested crude extracts of several plant species [sweet amber (*Hypericum androsaemum)*, fern (*Blechnum spicant*), nettle (*Urtica dioica*), moss (*Physcomitrium patens*), tobacco (*Nicotiana benthamiana*) and thale cress (*Arabidopsis thaliana*)] for their capability of providing protection to Vero E6 cells against the SARS-CoV-2 virus. As shown in Figure S2, certain degree of protection was observed for most of the tested plant species; the clearest protecting activity was observed for *Marchantia* crude extracts.

**Figure 2:**
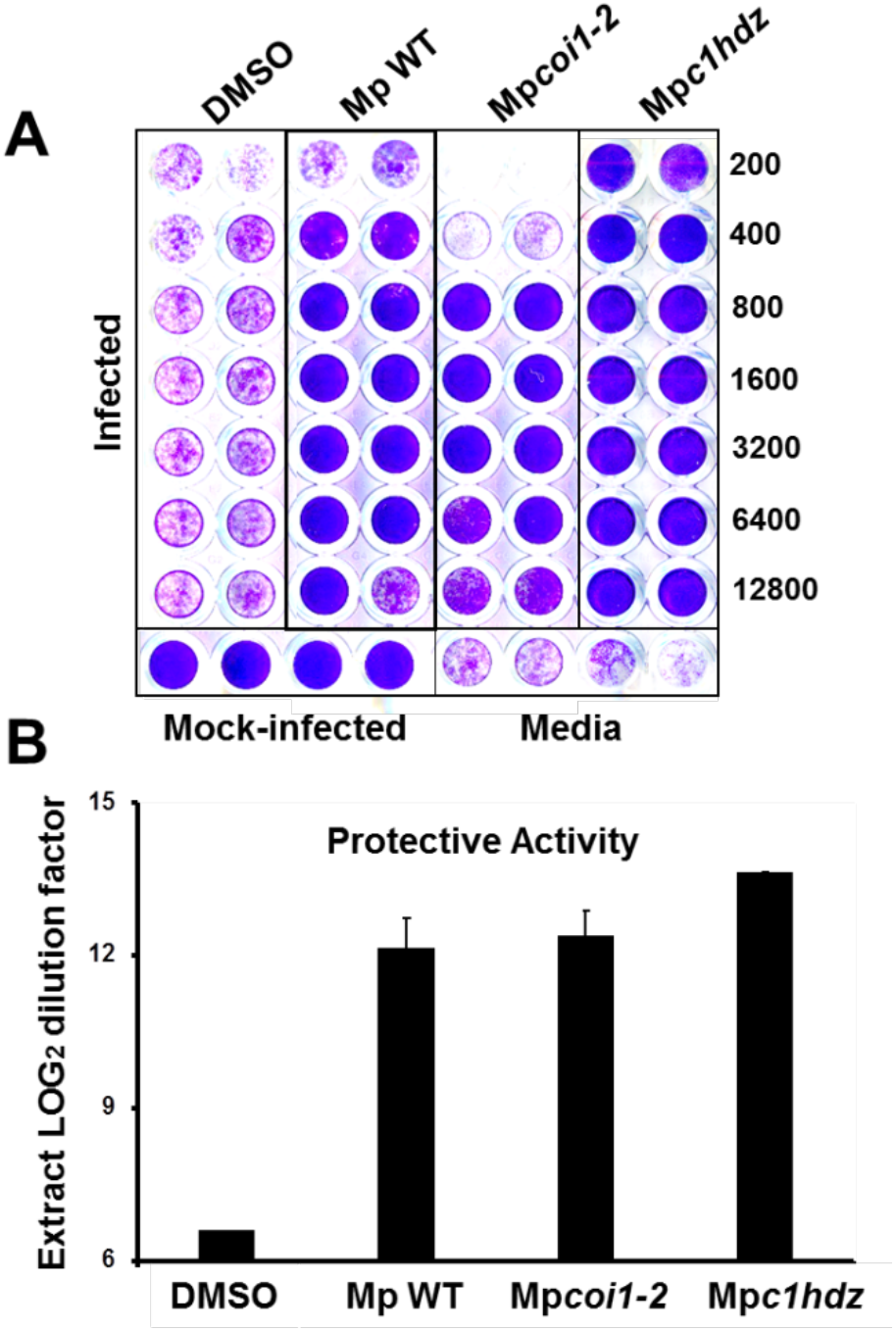
Antiviral candidates unlikely belong to Marchantia′s secondary metabolism. Vero E6 cells were inoculated with SARS-CoV-2 (MOI = 0.001) in the presence of serial 2-fold dilutions of crude extracts from WT, Mpcoi1-2 or Mpc1hdz Marchantia plants. Cultures were incubated for 72 h, time after which they were fixed and stained with a crystal violet solution. Mock-infected cells were used as the control of the integrity of the cell monolayer. A) Representative experiment showing dose-dependent protection of plant extracts in comparison with vehicle (DMSO)-treated cells. B) Numeric expression of the protective capacity as the log2 value of the highest dilution factor capable of full monolayer protection. Data are shown as average (± mean error) of two biological replicates.

### Identification of the antiviral metabolite

In order to identify the bioactive metabolite(s), we followed a bioactivity-guided chromatographic fractionation of the Marchantia’s WT crude extracts, which showed strong antiviral effect in previous assays. Chromatographic fractions were obtained via flash column employing a solvent polarity gradient, starting at *n*-hexane (100%) up to AcOEt:MeOH (4:1, v/v). A total of 56 fractions were obtained; fractions of a similar composition, based on their thin layer chromatography (TLCs) profiles, were combined and evaluated as 12 new pooled fractions (1-12). Fractions 10, 11 and 12 showed antiviral activity in the monolayer protection assay (Figure 3A-B). To directly confirm their antiviral activity, the viral antigen load reduction after inoculation of cell cultures with SARS-CoV-2 (MOI = 0.01) was measured. In this experimental setup, viral antigen accumulates as a consequence of virus propagation and can be quantitated using automated immunofluorescence microscopy. Figure 3C shows a dose-dependent reduction of viral antigen accumulation and the absence of cytotoxicity, as confirmed by normal cell numbers, estimated by DAPI staining and image analysis, and cell viability studies performed in parallel, uninfected cultures by MTT assays (Figure 3C).

**Figure 3:**
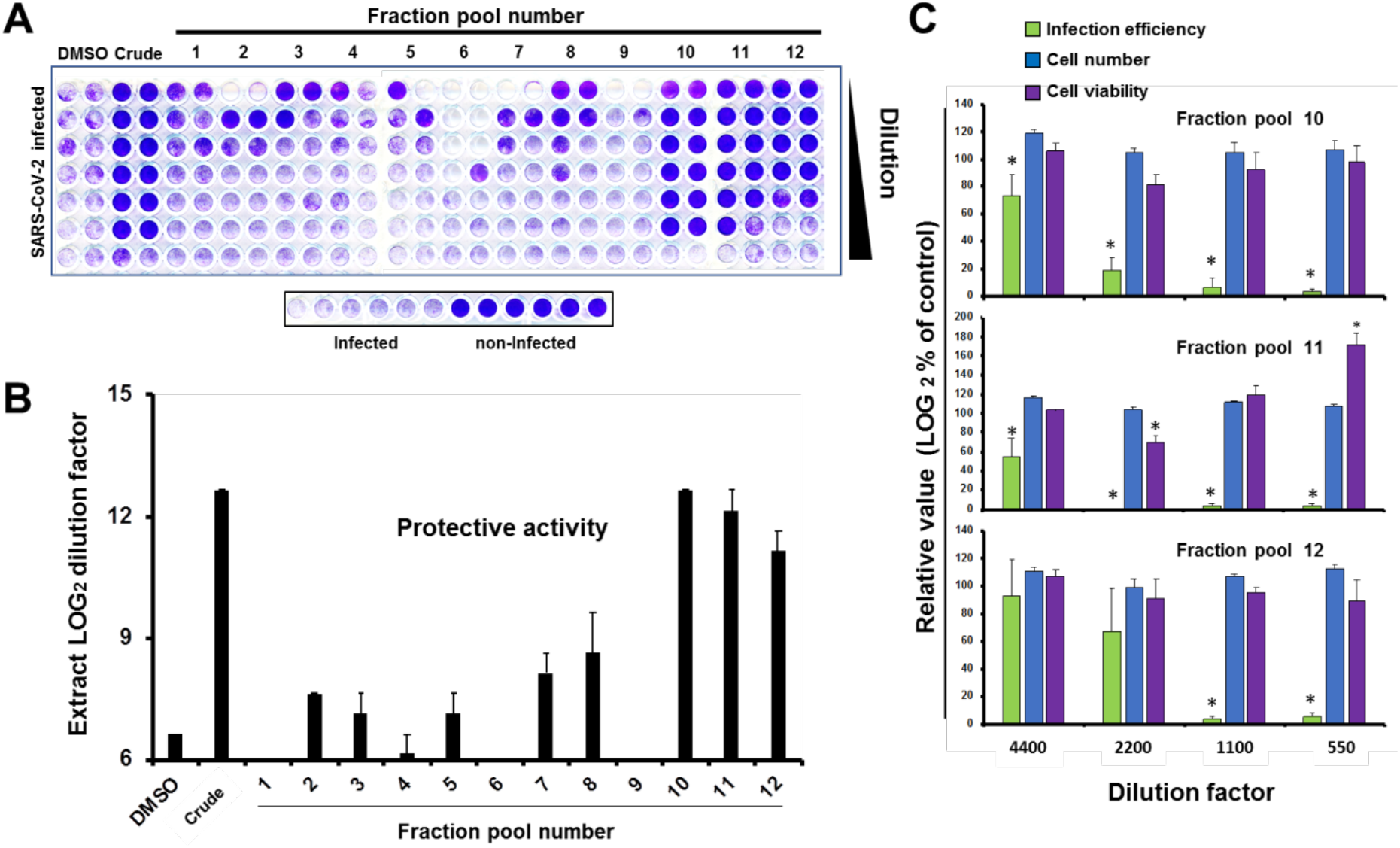
Extract fractionation and identification of antiviral fraction pools. Vero E6 cells were inoculated with SARS-CoV-2 (MOI = 0.001) in the presence of serial 2-fold dilutions of vehicle (DMSO), a crude Marchantia and the fraction pools. Inoculated cultures were incubated for 72 h, time after which they were fixed and stained with a crystal violet solution. Mock-infected cells were used as the control of the integrity of the cell monolayer (non-infected). A) Experimental plates showing cell monolayer integrity (purple) in the presence of fractions containing the antiviral compound(s). B) Numeric expression of the protective capacity as the log2 value of the highest dilution factor capable of full monolayer protection. Data are shown as average and mean error of two biological replicates. C) Vero E6 cells were inoculated (MOI = 0.01) in the presence of the indicated fraction dilutions and incubated for 24 h before fixation and processing for immunofluorescence microscopy and cytotoxicity assays as described in the materials and methods section. Data are shown as average (± SD) of three biological replicates. Statistical significance was estimated using one-way ANOVA (Dunnet′s post-hoc test, *p<0.05).

Interestingly, TLCs of the three active fractions presented red fluorescent spots (under long wave ultraviolet light, 365 nm; Supplementary Figure S3A), which are characteristic of plant chlorophylls, but with a smaller retention factor [Rf = 0.36, AcOEt:MeOH (9:1, v/v)] than that of chlorophyll (Rf = 0.92). At this point, we suspected that the active antiviral metabolite(s) could be related to plant chlorophylls, especially because a weak antiviral activity was observed at low dilutions of fractions 2 and 3 (Figure 3A), both containing chlorophyll. Indeed, these fractions, when re-chromatographed, yielded spots on the TLC with an Rf consistent with that observed in the active fractions 10 to 12, indicating that the active metabolite is a chlorophyll derivative (Supplementary Figure S3B). It is worth mentioning that heat notably accelerated the chlorophyll decomposition into the investigated metabolite.

Next, we employed preparative TLC in order to better isolate and characterize the red-light emitting metabolite. Photosynthetic metabolites were extracted from *M. polymorpha* WT and Mp*c1hdz* plants, and the extracts subjected to preparative TLC. Three spots showed fluorescence in the vicinity of the expected Rf; these spots where isolated, analysed by HPLC-UV-MS and the bioactivity assayed as fractions C, D and E (Figure 4A). Fraction D showed the strongest antiviral activity (Figure 4B), which corresponded with an enrichment of compound **1** in the UV chromatograms (Figure 4C). Careful analysis of the MS spectra (positive mode) revealed a molecular formula of C_35_H_36_N_4_O_5_ for compound **1**, as deduced by HR-ESI^+^-MS from its monoprotonated molecular ion, [M+H]^+^, with a *m/z* of 593.2759. The identified molecular formula (containing four nitrogen atoms), the exact mass and the characteristic red fluorescence of the compound helped to identify **1** as the chlorophyll catabolite *Pheophorbide a* (PheoA, Figure 4C). The identity of PheoA was further confirmed by comparison with both a commercially available original standard (Santa Cruz Biotechnology) and a semisynthetic sample. PheoA was obtained semi-synthetically from *Marchantia* in good overall yield via a solvent-free, thus, environmentally friendly method (Supplementary Figure S4 and M&M section).

**Figure 4:**
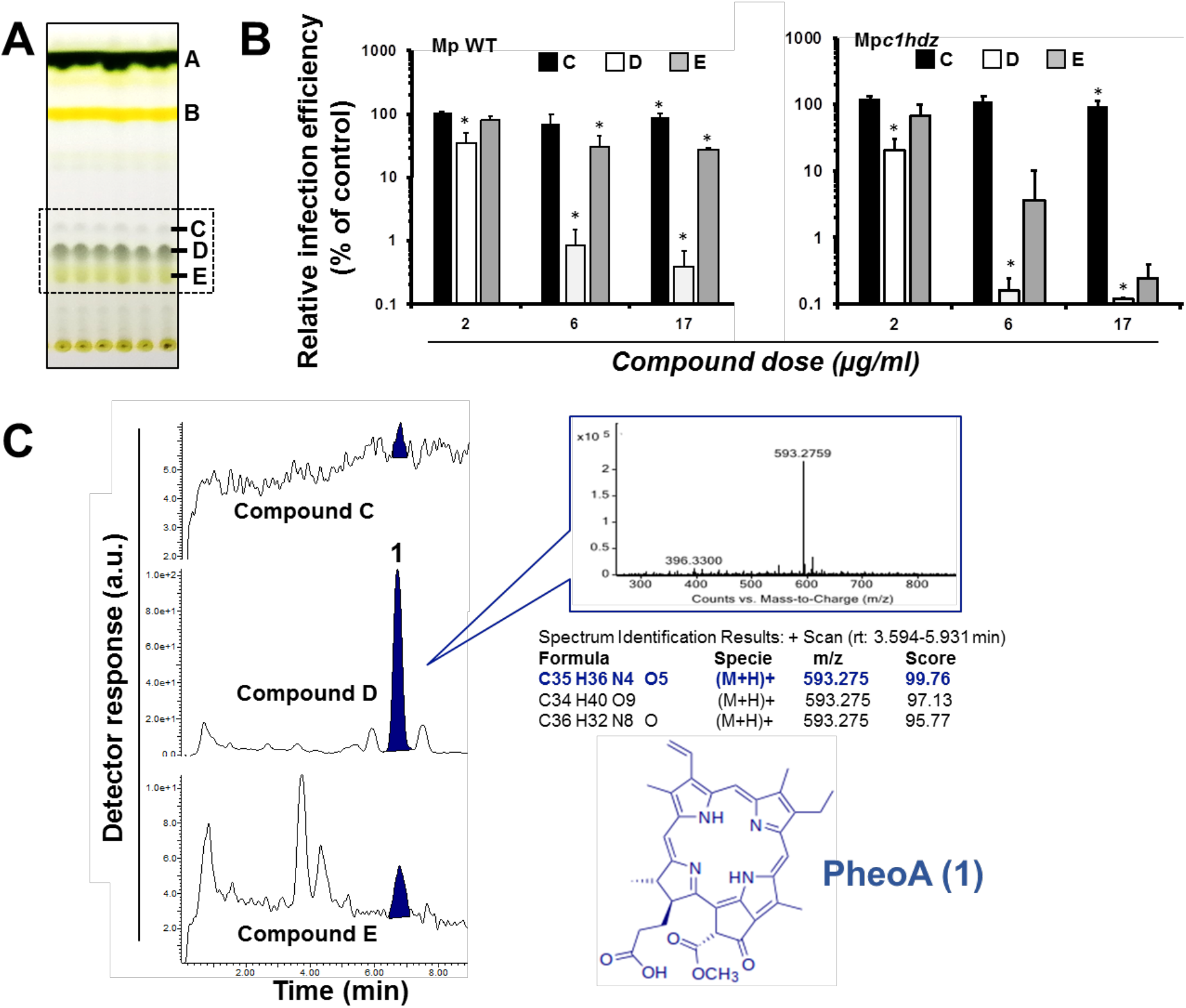
Identification of the main antiviral compound in Marchantia extracts. A) Representative TLC of WT and Mpc1hdz Marchantia extracts. Compounds C, D and E (box) were tested for their antiviral potential. B) Vero E6 cells were inoculated (MOI = 0.01) in the presence of the indicated compounds and incubated for 24 h before fixation and processing for immunofluorescence microscopy. Data are shown as average and SD of three biological replicates. Statistical significance was estimated using one-way ANOVA and a Dunnet′s post-hoc test (*p<0.05). C) Representative HPLC/MS analysis (shown for WT) of fractions C, D, and E, including exact mass determination of the antiviral candidate **1,**and its inferred chemical structure.

#### Antiviral activity of PheoA

To confirm the antiviral potential of PheoA, a commercially available PheoA stock solution was serially diluted and mixed with a virus stock to inoculate Vero E6 and Huh7-ACE2 cells (human hepatoma cells expressing ACE2). Cells were fixed 72 h post-inoculation and stained with crystal violet to visualize the integrity of the cell monolayer. Figure S5 shows consistent protective capacity of PheoA at concentrations above 40 ng/ml (67 nM) in both cell lines.

PheoA antiviral activity was further confirmed by immunofluorescence microscopy, to estimate virus propagation, and an MTT assay to evaluate compound cytotoxicity. PheoA dose-response curves demonstrate PheoA’s antiviral activity against SARS-CoV-2 in Vero E6 and human lung epithelial cells (A549-ACE2 and Calu3); in all three models, no cytotoxicity was observed (Figure 5A-C). This dataset was used to determine effective concentrations (EC_50_ and EC_90_) and cytotoxicity indexes. The estimated EC_50_ and EC_90_ values were around 14 ng/mL (25 nM) and 156 ng/mL (86 nM), with a wide therapeutic window in all tested cell lines (Table 1).

**Figure 5:**
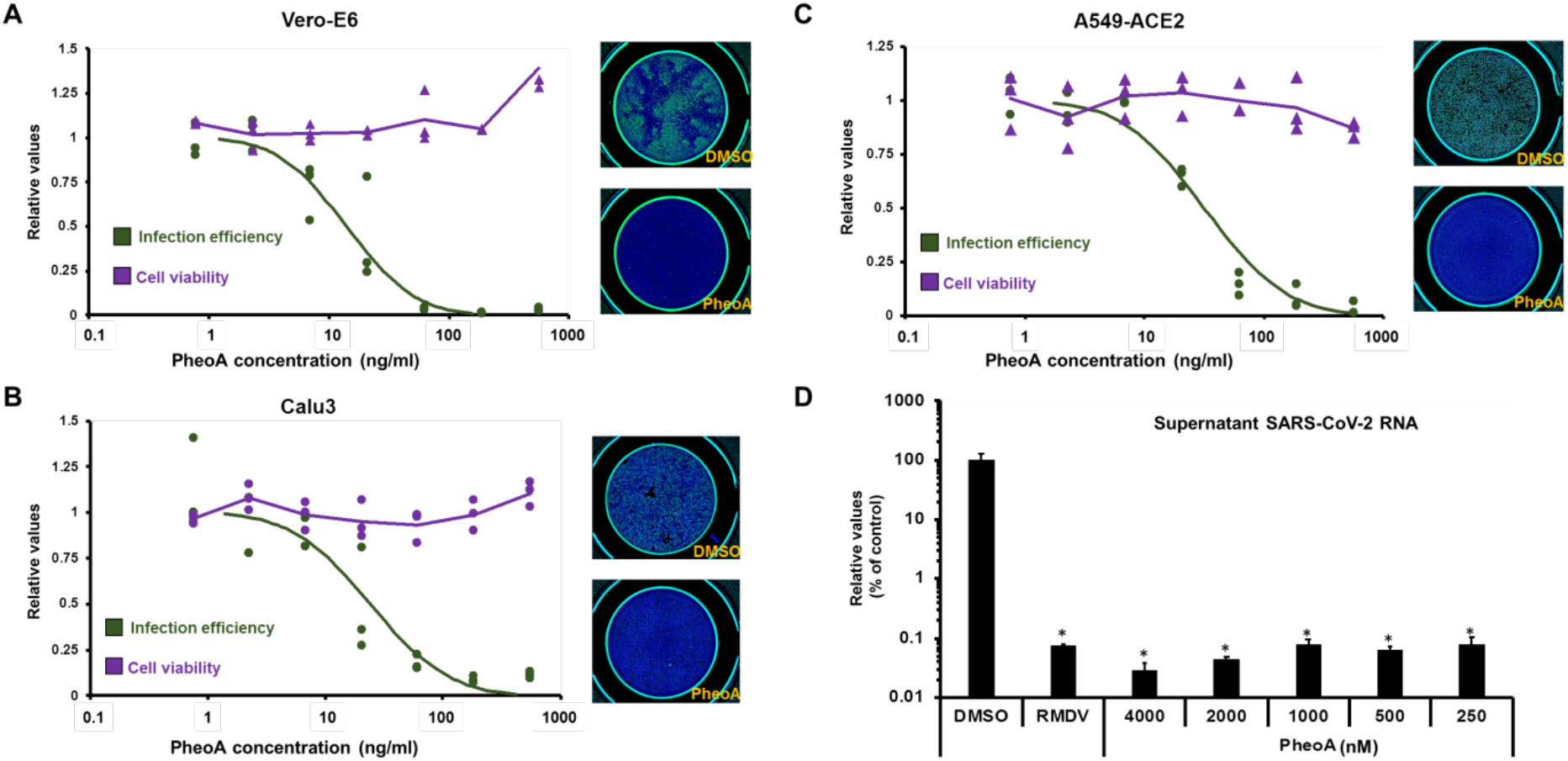
Pheophorbide a shows antiviral activity against SARS-CoV-2 in Vero E6 cells and human lung epithelial A549-ACE2 and Calu3 cell lines. Commercially available PheoA was serially diluted and mixed (1:1) with SARS-CoV-2 preparations to achieve the indicated compound concentrations and a final MOI of 0.005 for (A) Vero E6 and (B) Calu3 and 0.01 for (C) A549-ACE2 cells. Cultures were incubated for 48 h, fixed and processed for automated immunofluorescence microscopy analysis. Parallel, uninfected cultures were processed for cytotoxicity evaluation using an MTT assay. Relative infection efficiency data (N=3 per dose) are shown as individual data and a PROBIT regression curve (green line) using the represented values. Cytotoxicity data (N=3 per dose) are shown as the individual data and a moving average trendline. D) A549-ACE2 cells were inoculated at MOI = 0.01 in the presence of increasing concentrations of PheoA or 5000 nM remdesivir and incubated for 48 h. Samples of the supernatants were collected, heat-inactivated and directly subjected to RT-qPCR to estimate overall infection efficiency. Data are expressed as relative values compared with the vehicle (DMSO)-treated cells and are shown as mean and standard deviation of three biological replicates (N=3). Statistical significance was estimated using one-way ANOVA and a Dunnet′s post-hoc test (*p<0.05).

**Table 1:** Potency and cytotoxicity indexes of commercially available PheoA

| Cell line | EC<sub>50</sub> (nM) | EC<sub>90</sub> (nM) | CC<sub>50</sub> (nM) |
| --- | --- | --- | --- |
| Vero-E6 (green monkey) | 25 | 86 | >8420 |
| A549-ACE2 (human lung) | 52 | 263 | 2020 |
| Calu3 (human lung) | 34 | 232 | >8420 |

To fully evaluate the extent to which PheoA interferes with SARS-CoV-2 infection (i.e. on viral replication) - given the narrow dynamic range of immunofluorescence -, we employed RT qPCR to determine the extracellular viral RNA load in human cells inoculated with SARS CoV-2 (MOI = 0.001). In this experimental setup, remdesivir (5 μM) reduced extracellular viral RNA by three orders of magnitude. Interestingly, PheoA significantly reduced viral spread even at the lowest tested concentration (150 ng/mL; 0.25 μM), as shown by the three times log reduction in viral RNA accumulation after a 48h incubation period (Figure 5D). These results indicate that PheoA displays a remarkable potency and a satisfactory therapeutic index, and suggest that it may be considered as a potential candidate for antiviral therapy against SARS-CoV-2. Furthermore, results also suggest that PheoA is a major determinant of the antiviral activity observed in crude *Marchantia* extracts (Figure 1). This notion is underscored by the fact that the semisynthetic PheoA (88-94% by HPLC-UV/Vis) showed comparable potency to crude extracts in the different cell lines (Supplementary Figure S6). Nevertheless, other related chlorophyll metabolites may also contribute with antiviral activity. In fact, pyropheophorbide a (pPheoA), which was also found in antiviral fractions was tested to verify its antiviral potential in Vero E6 cells. The pPheoA showed antiviral activity in the absence of measurable cytotoxicity with an EC_50_ of 185 nM (Supplementary Figure S7), suggesting lower potency than PheoA and further underscoring a major role for PheoA in the antiviral activity of Marchantia extracts.

#### Antiviral spectrum of Pheophorbide A

PheoA has previously been proven as an antiviral against the hepatitis C virus (HCV)^27^ and virucidal against herpes simplex virus (HSV)^28^. Thus, we determined the antiviral spectrum of PheoA on different enveloped +RNA viruses. First, we confirmed antiviral activity against HCV, using a surrogate model of infection by propagation-deficient, *bona fide* reporter virus bearing a luciferase reporter gene generated by trans-complementation (HCVtcp). Dose-response curves of the luciferase activity in HCVtcp-infected Huh7 cells indicated an EC50 of 177 ng/mL (300 nM) for PheoA against HCV (Figure 6A), very similar to the previously reported EC_50_^27^.

Next, we asked whether PheoA antiviral activity against SARS-CoV-2 could also be observed against other human coronaviruses such as hCoV-229E (Genus alphacoronavirus; Subgenus Duvinecovirus), which has been associated with mild respiratory infections^29^. Huh7 cells were inoculated inoculated with a GFP reporter-expressing recombinant hCoV-229E (MOI = 0.01) and total GFP expression in the target Huh7 cells was evaluated by automated microscopy 48 h post-inoculation. Similar to the results with the HCV infection model, PheoA reduced viral spread (EC_50_ of 76 ng/mL; 128 nM) while no cytotoxicity was observed (Figure 6B). Strikingly, comparable results were obtained in an experimental model of infection by the West Nile Virus, a mosquito-borne zoonotic pathogen that may cause encephalitis in infected humans. A recombinant, di-cistronic infectious molecular clone expressing GFP in the second cistron^30^ was used to inoculate Huh7 cells (MOI = 0.01). Cells were imaged 48 h post infection and the degree of virus propagation was determined via automated microscopy; dose response curves (Figure 6C) indicated antiviral activity for PheoA with an EC_50_ of 38 ng/mL (68 nM). Altogether, these results suggest that PheoA displays a broad-spectrum antiviral activity against +RNA viral pathogens. In view of these results, we decided to test PheoA antiviral activity against the vesicular stomatitis virus (VSV), a negative-strand RNA (-RNA) virus. Dose-response experiments, similar to those describe above, were performed employing A549-ACE2 cells. Interestingly, the VSV-GFP^31^ was not susceptible to PheoA doses largely exceeding those for which antiviral activity was observed against SARS-CoV-2 (Figure 6D). Collectively, these results suggest that PheoA is a broad-spectrum antiviral and that +RNA viruses are particularly susceptible.

**Figure 6:**
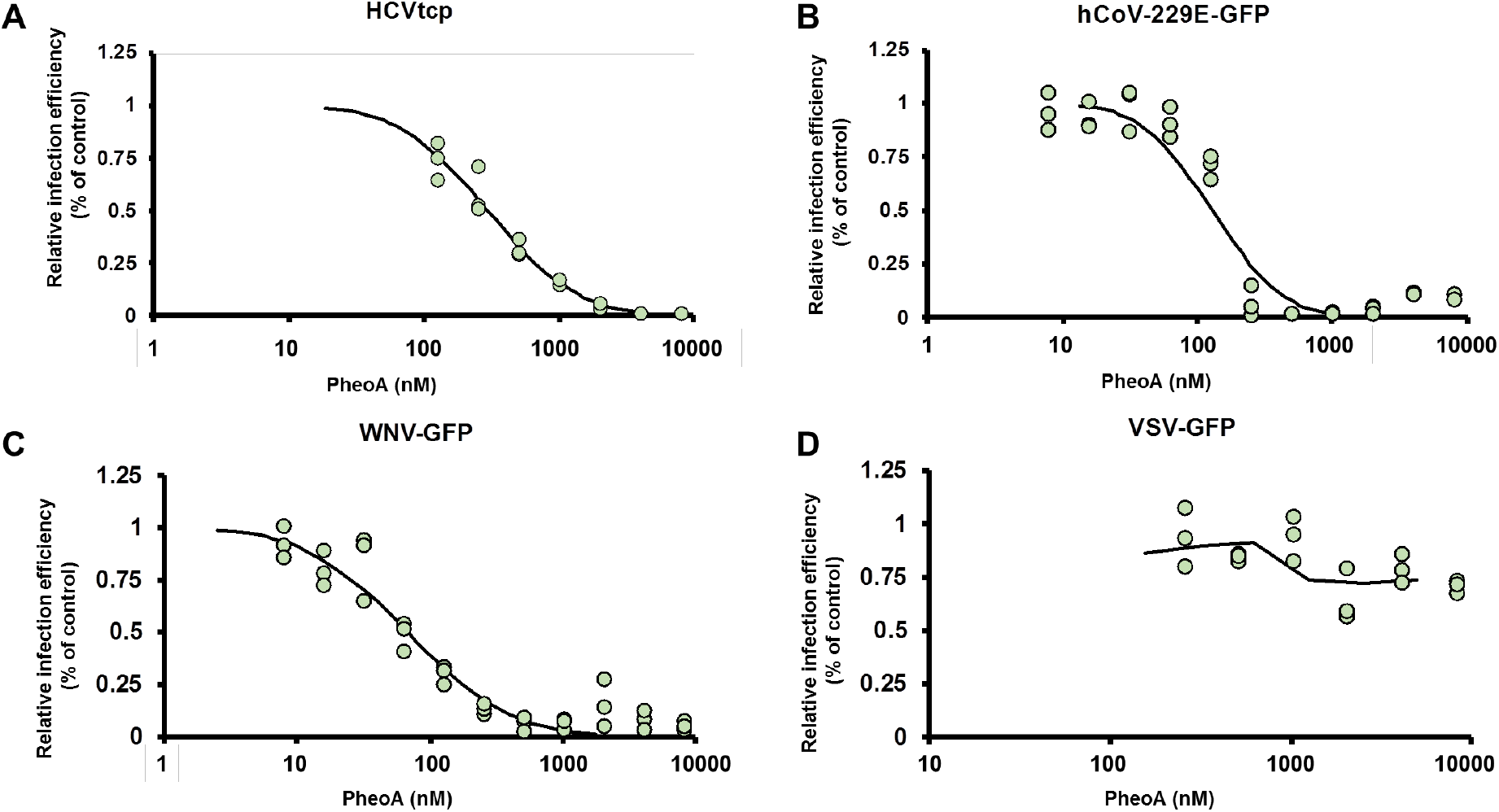
Antiviral spectrum of PheoA against different RNA viruses. The effectiveness of PheoA was tested against four different recombinant RNA viruses expressing reporter genes. A) Huh7 cells were infected with HCVtcp in the presence of increasing PheoA doses and luciferase activity was determined 48 h post-inoculation. (B-D) Cells were inoculated in the presence of increasing concentrations of PheoA at MOI 0.01 and incubated to enable virus propagation. At the endpoint, cells were fixed and counter-stained with DAPI to control for unexpected cytotoxic effects. Relative infection efficiency was estimated using automated microscopy and is expressed as percentage of the infection efficiency observed in control wells. B) Huh7 cells were infected with hCoV-229E-GFP and fixed 48 h post-inoculation. C) Huh7 cells were infected with WNV-GFP and fixed 48 h post-inoculation. D) A549-ACE2 cells were inoculated with VSV-GFP and fixed 16 h post-inoculation. Individual replicate data are shown as green dots (N=3) and the PROBIT regression curve used to estimate EC50 values is shown.

#### Pheophorbide a can be employed in combination with remdesivir

Once the broad antiviral activity of PheoA has been demonstrated, we studied whether the addition of PheoA to remdesivir treatment could result in a synergistic effect on viral infection. Thus, combination treatments were performed with increasing doses of PheoA and remdesivir. Drugs were mixed in different proportions, combined with infectious SARS-CoV-2 (MOI = 0.01) and the mixtures were used to inoculate Vero E6 cells. Twenty-four hours later, cells were fixed and processed to determine the infection efficiency as described in Figure 5. Individual treatment with either compound resulted in the expected dose-dependent inhibition of virus infection, achieving the EC50 at the expected doses (2000 nM for remdesivir and 40 nM for PheoA). Increasing concentrations of PheoA improved remdesivir efficacy and viceversa. However, full analysis of the combinations resulted in a synergy index close to three, indicating that the drug combination is mostly additive^32^, with an area of synergy at concentrations close to the EC50s (Figure 7). These results suggest that combinations of PheoA with other antivirals may result beneficial, as it was observed by its additive effect in combination with remdesivir in cell culture infection models.

**Figure 7:**
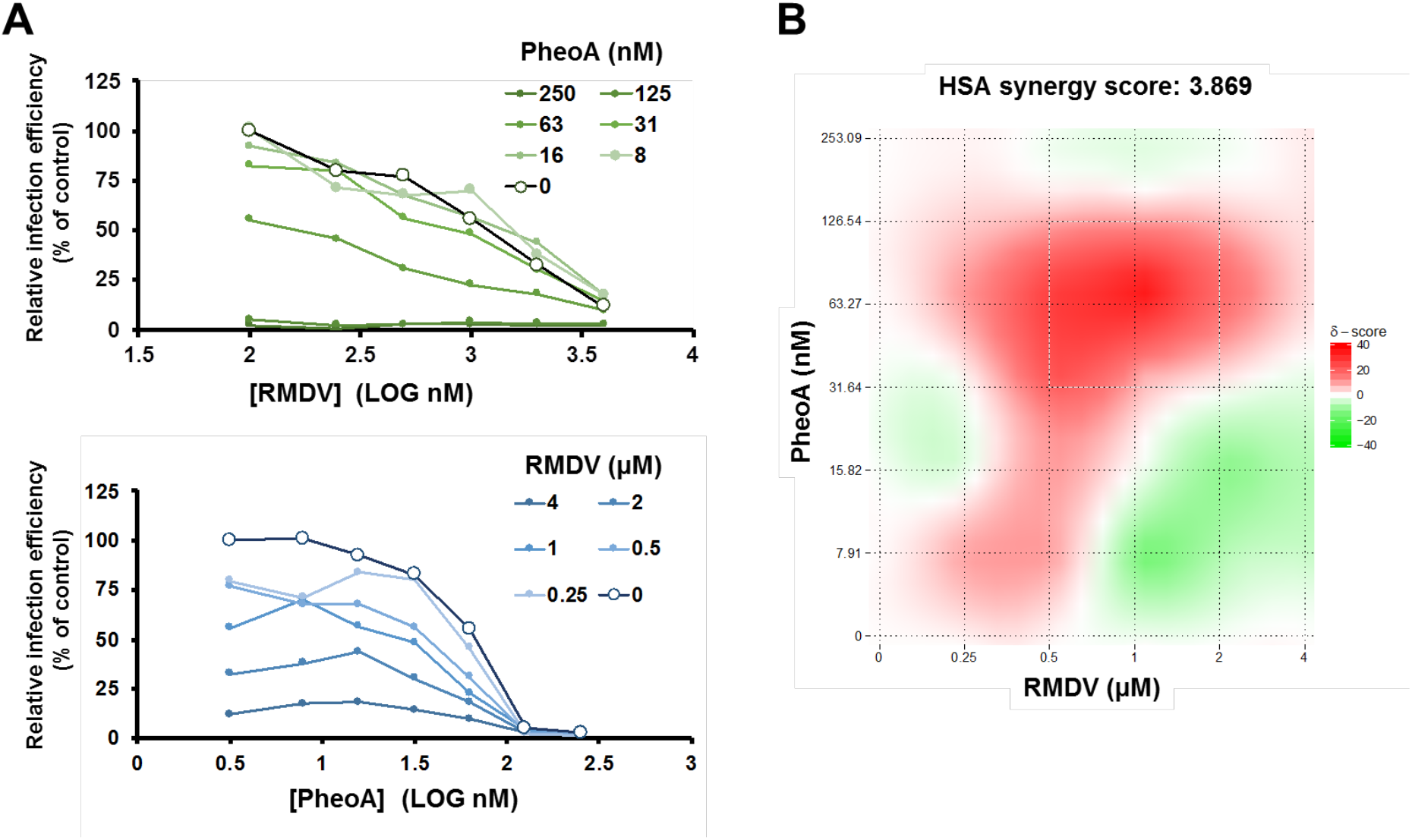
Combination treatment of Pheophorbide a with remdesivir. Vero E6 cells were inoculated at MOI = 0.005 in the presence of increasing concentrations of PheoA in combination with increasing doses of remdesivir. Twenty-four hours post infection, cells were fixed and processed for automated immunofluorescence microscopy. Relative infection efficiency values were estimated as percentage of the values obtained in mock-treated cells. A) Data are shown as average of two biological replicates. B) Heatmap describing the areas of synergy within the combination treatments.

### Characterization of Pheophorbide a mode of action on SARS-CoV-2 infection

Next, antiviral efficacy of PheoA was compared when PheoA was (i) present at all times, (ii) added only during virus inoculation, or (iii) added only after virions had effectively penetrated the cells in single-cycle infection experiments (MOI = 5). Imatinib (15 μM), for which antiviral activity at the level of virus entry has previously been demonstrated^33^, was employed as the reference compound. Infection efficiency revealed the expected antiviral activity for imatinib and PheoA when maintained at all times in the experiment (Figure 8A). Imatinib showed comparable efficacy when added during the virus entry phase and greatly lost efficacy when added after virion internalization, as expected for an entry inhibitor. Similar results were obtained with PheoA, inhibition was nearly identical when maintained at all times or added during viral entry, but only ca. 6% of the maximum efficacy was observed when added after virion internalization. These results suggest that PheoA is mainly acting at early stages of the infection, potentially at the level of viral entry.

**Figure 8:**
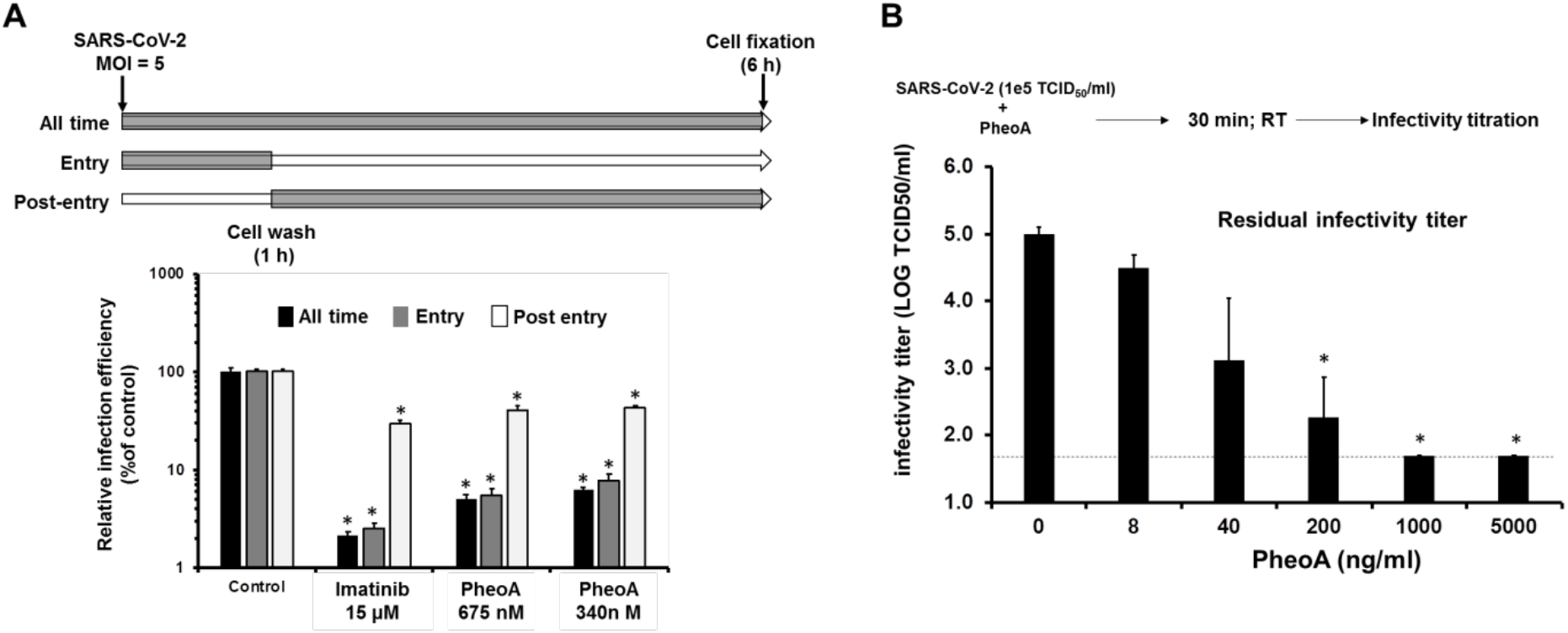
PheoA shows virucidal activity against SARS-CoV-2 interfering with early aspects of the infection. A) *Time-of-addition experiments indicate that PheoA interferes with early aspects of SARS-CoV-2 infection.* Vero E6 cells were inoculated at MOI = 5 in the presence (gray) or absence (white) of the indicated doses of PheoA or imatinib as described in both the text and the scheme. Cells were incubated for 6 h in the presence (gray) or absence (white) before chemical fixation and processing for immunofluorescence microscopy. Infection efficiency is expressed as the percentage of that observed in vehicle DMSO-treated cells and is shown as average and standard deviation of three biological replicates (N=3). Statistical significance was estimated using one-way ANOVA and a Dunnet′s post-hoc test (*p<0.05). B) *Pheophorbide A shows virucidal activity against SARS-CoV-2.* SARS-CoV-2 virus stocks were diluted to obtain 1*10^5^ TCID50/mL and were mixed with increasing concentrations of PheoA or the vehicle (DMSO). Virus-compound mixtures were incubated at room temperature for 30 minutes and were serially diluted to determine the remaining infectivity titer using endpoint dilution and determination of virus-induced cytopathic effect by crystal violet staining in Vero-E6 cells. Values are expressed as LOG TCID50/mL and shown as the average and standard deviation of three independent experiments (N=3). Statistical significance was estimated using one-way ANOVA and a Dunnet ́s post-hoc test (*p<0.05).

In view of these results, we directly tested this hypothesis by determining the antiviral activity of PheoA in a surrogate model of infection recapitulating only aspects related with viral entry such as receptor recognition, virion internalization or membrane fusion. This system is based on reporter retroviral vectors pseudotyped with SARS-CoV-2 Spike protein (Spp) or VSV glycoprotein as a control (VSVpp). Infection efficiency in the presence of antiviral molecules is determined as the relative expression values of the reporter gene, in this case a Firefly luciferase^33^. Relative infection efficiency was measured in the presence of the entry inhibitor imatinib (15 μM) and antiviral doses of PheoA. As expected, imatinib selectively inhibited Spp and not VSVpp entry (Supplemantary Figure S8). PheoA barely interfered with either retroviral pseudotype infection efficiency, with a maximum reduction of 40% at the maximum dose (400 ng/ml; 678 nM) (Supplementarty Figure 8). These results suggest that, while time-of-addition experiments suggest that PheoA interferes predominantly with early aspects of the infection, surrogate models of viral entry indicate that it does not interfere substantially with molecular events leading to viral entry *per se*.

PheoA irreversibly interferes with the virion infectivity in HSV and influenza infection models^28^, therefore, we explored whether PheoA could be virucidal for SARS-CoV-2 virions, a property that would be compatible with the observations made in the time-of-addition experiments. Thus, a known number of infectious particles (10^5^ TCID_50_) were mixed with increasing doses of PheoA [8 ng/mL(0.014 μM) to 5000 ng/mL (8.45 μM)] and incubated for 30 minutes before residual infectivity titer was calculated by endpoint dilution and TCID_50_ determination. The dose of PheoA was kept below its effective concentrations during the titration experiment. Figure 8B shows how pre-incubation of the infectious virions with PheoA (40 ng/mL or more) resulted in irreversible, dose-dependent inactivation of the virus infectivity. These results suggest that PheoA is virucidal for SARS-CoV-2 virions and that virion infectivity inactivation contributes to its overall antiviral effect.

## Discussion

Due to their metabolic richness plants have been traditionally used as source of medicines. The potential of plant metabolites in pharmacology is still far from being saturated, particularly in certain plant clades. Indeed, bryophytes (non-vascular plants) are particularly rich in specialized metabolites that are rarely found in other plant lineages^34^. Here, we explored this richness in order to find antiviral compounds against the SARS-CoV-2 virus by employing an activity-guided chromatographic method; and identified PheoA as a potent antiviral, very efficient not only against SARS-CoV-2 but also against several other enveloped viruses.

The first evidence for antiviral activity of PheoA derives from observations made on HSV infection models^35^. In those initial reports, some degree of selectivity towards other viruses was reported, since adenovirus (Type VI), Japanese Encephalitis virus (JEV) or poliovirus were not affected by treatment with PheoA-enriched algal extracts^36^. Subsequent studies suggested that PheoA and pPheoA display broad-spectrum antiviral activity against enveloped viruses, including influenza A^28^ and HIV^37^, but not against non-enveloped viruses^28.^ Ohta et al. reported that PheoA-containing preparations may display virucidal activity against HSV^36^, a concept that was further supported by Bouslanu et al.^28^. Our observations support that PheoA inactivates SARS-CoV-2 (enveloped +RNA virus) virion infectivity through a virucidal mode of action. First, time-of-addition experiments indicate that early aspects of the infection are targeted by PheoA. Second, the study of viral entry using retroviral pseudotypes did not reveal any measurable antiviral activity against SARS-CoV-2 entry, suggesting that receptor recognition by the Spike protein, particle internalization and Spike-mediated fusion are not affected by PheoA. Similar models have been used to identify key SARS-CoV-2 entry factors as well as to study antibody neutralizing activity^38,39^. Thus, irreversible inactivation of viral infectivity (virucidal activity) was tested as a possible mechanism reconciling these apparently contradicting observations. Pre-incubation of infectious SARS-CoV-2 virions with PheoA rendered the virions non-infectious even when PheoA-virus dilutions were performed below active PheoA concentrations. These observations are similar to those reported by other groups in other infection models^28,36^. The virtual lack of activity of PheoA at post-entry levels may be explained by the fact that PheoA can only act on the viral particle, or that PheoA requires longer incubation periods to penetrate the cell and interfere with downstream steps of the virus lifecycle. Given that overall effectiveness of PheoA as virucidal is substantially stronger than during multiple cycle infection experiments, it is likely that virucidal activity is the main mechanism for interference with SARS-CoV-2 infection.

PheoA has been shown to integrate into biological membranes^40^. Thus, it is possible that PheoA could insert into the viral envelope lipid bilayer, altering its biophysical properties, or even disrupting it, thusrendering the virion non-infectious. This would explain PheoA’s virucidal activity and its broad-spectrum among enveloped viruses. However, some degree of selectivity was observed since PheoA doses that completely abolished infection by several +RNA viruses did not interfere with VSV or retroviral pseudotype (both also enveloped) infection. It likely is possible that the membrane’s lipidic composition could play a key role for PheoA incorporation. In this sense, VSV and retroviral pseudotypes are assembled at the plasma membrane^41,42^, while the rest of the tested virus particles are assembled in intracellular compartments^43,44^, which display a very different membrane composition from that of the plasma membrane^45^. PheoA is a plant derived porphyrin closely related to animal porphyrins, which have been widely described as broad-spectrum virucidals (reviewed in Sh. Lebedeva et al.^46^). Virion inactivation is thought to occur through incorporation of porphyrins into the viral envelope membrane and modifying its physico-chemical properties, thus interfering with host cell recognition and fusion processes. However, porphyrins such as protoporphyrin IX display antiviral activity independently of their virucidal activity at post-entry steps and have been proposed to interfere with receptor (ACE2) recognition in SARS-CoV-2 infection models^19^. The structural resemblance between PheoA and proporphyrin IX may explain their similar antiviral properties (broad spectrum and virucidal), but, at the same time, their differences may contribute to PheoA’s increased tolerability and *in vivo* effectiveness, an issue that has extensively been explored for different PheoA applications as photosensitizer in photodynamic therapies against cancer ^47,48^.

One huge advantage of PheoA is that it is readily available from plant and algae chlorophyll. PheoA is the dephytylation and demetallation product of chlorophyll *a*, processes mediated by chlorophyllase and Mg-dechelatase, respectively ^49,50^. Clorophyllase activity is favored by high temperatures (60-80 °C)^49^ and its accumulation varies throughout plant development and in stress conditions. In this study, we also exploited stress conditions (heat) that favour PheoA accumulation and PheoA was semisynthetically prepared from *M. polymorpha* in good overall yield.

Another advantage of PheoA is that its combination with remdesivir has an additive effect with no cross inhibition in their antiviral activity, and a mild synergy. This, together with its low toxicity *in vivo,* represents an advantage that could be clinically exploited^51^.

## Materials and Methods

### Equipment and reagents

All solvents were of ACS quality unless stated otherwise. Commercially available PheoA was purchased from Santacruz Biotechnology (>90% by HPLC). A Geno Grinder Spex/SamplePrep 2010 was employed for tissue homogenization. Glass or aluminium supported Silica gel 60 (Merck) was used for preparative and analytical TLCs, respectively; for flash column purification, silica gel 60 □, 230-400 mesh, 40-63 μm was employed. HPLC-UV-MS analysis was carried out by using a Waters Separations module Alliance e2695 system, a Waters QDa Detector Acquity QDa and a Waters Photodiode Array Detector 2996. HPLC was performed by using HPLC grade solvents and a Sunfire C18 (4.6 × 50 mm, 3.5 μm) column at 30 °C, with a flow rate of 1 mL/min and a mobile phase gradient from 70 to 95 of A (formic acid 0.1% in CH_3_CN) in B (0.1% of formic acid in H_2_O) for 10 minutes. Electrospray in positive mode was used for ionization. The HR-MS analysis was carried out by using an Agilent 1200 Series LC system (equipped with a binary pump, an autosampler, and a column oven) coupled to a 6520 quadrupole-time of flight (QTOF) mass spectrometer. CH_3_CN:H_2_O (75:25, v/v) was used as the mobile phase at 0.2 mL/min. The ionization source was an ESI interface working in the positive-ion mode. The electrospray voltage was set at 4.5 kV, the fragmentor voltage at 150 V and the drying gas temperature at 300 °C. Nitrogen (99.5% purity) was used as nebulizer (207 kPa) and drying gas (6 L/min). The scan range was 50–1100 m/z.

### Preparation of crude M. polymorpha extracts

Plant material (10 g, fresh weight) was collected and dried in an oven (60 °C) until constant weight. The dry tissue was ground to a fine powder using a Geno/Grinder (2x 2 min at 2700 rpm) and extracted two times at room temperature with 30 mL of CHCl_3_:MeOH (2:1, v/v) for at least 6 h each time. Extracts were combined and concentrated under a nitrogen flow. The remaining solid was dissolved in DMSO (1 mL) to create the stock solutions employed in the bioassays.

### Chromatographic fractionation of extracts

Plants extracts were prepared as described above starting from ca. 20 g of plant material and directly subjected to silica gel flash column chromatography employing a solvent polarity gradient starting at *n*-hexane (100%) up to AcOEt:MeOH (4:1, v/v). A total of 56 metabolite-enriched fractions were obtained. Fractions were analyzed by TLC and those of similar composition were combined to render 12 new pooled fractions, which were screened for antiviral activity.

### Preparative TLCs

Photosynthetic metabolites were extracted [two times, o/n, acetone (30 mL)] from fresh, finely grounded *M. polymorpha* thallus (ca. 40 g). The combined extracts were concentrated to a final volume of 10 mL, centrifuged (4000 rcf), filtrated (45 μm, Whatmann PTFE filters) and chromatographed on preparative TLC plates employing the solvent system AcOEt:MeOH (9:1, v/v).^1^ Selected fluorescent spots (C, D and E) were scraped off, eluted (AcOEt:MeOH, 4:1, v/v) and dried under a nitrogen stream. Single components (C, D and E) were prepared at a 10 mg/mL and submitted to both antiviral assays, as described below, and HPLC-UV-MS analysis as described above.

### Semisynthetic preparation of PheoA

Fresh *M. polymorpha* thallus (ca. 50 g) was oven dried to produce ca. 3.5 g of dry material, which was ground to a fine powder in the GinoGrinder as described above. The obtained powder (3 g) was extracted (3x, Acetone, 90 mL), the extracts combined and silica gel (6 g, ratio 2:1 by weight relative to the dried leaf powder) added. The solvent was evaporated under reduced pressure to produce an impregnated silica, which was heated (60 °C) overnight to further potentiate PheoA production. The obtained silica was directly loaded into a flash column that was run as follows: column ID = 3 cm, silica (70 g), *n*-hexane:AcOEt (1:1, 300 mL), AcOEt:MeOH (9:1, 300 mL; 4:1, 600 mL; 7:3, 300 mL), fractions of ca. 35 mL were collected. Fractions were analyzed by TLC and the obtained PheoA (3.9 mg, 0.13 % from oven dried leaf material).

### Cell culture

Vero E6 (ATCC) and Calu3 (ATCC) cell lines were kindly provided by Dr. Enjuanes (CNB-CSIC). A549 cells were kindly provided by Dr. Juan Ortín (CNB-CSIC) and Huh7 cells were kindly provided by Dr. Chisari (TSRI, La Jolla). A549 and Huh7 cells were transduced with a retroviral vector enabling expression of ACE2 in a di-cistronic expression cassette also conferring resistance to blasticidine. Transduced populations were selected using 2.5 μg/mL of blasticidine. All cell cultures were kept in complete media (DMEM) supplemented with 10 mM HEPES, 1X non-essential amino acids (gibco), 100 U/mL penicillin-streptomycin (GIBCO) and 10% fetal bovine serum (FBS; heat-inactivated at 56 °C for 30 min). Unless otherwise stated, all infection experiments were performed at 37oC in a CO_2_ incubator (5% CO_2_) the presence of 2% FBS and in the absence of selection antibiotics.

### Viruses

SARS-CoV-2 (Orthocoronavirinae; Alphacoronavirus; Sarbecovirus; strain NL/2020) was kindly provided by Dr. R. Molenkamp, Erasmus University Medical Center Rotterdam. SARS-CoV2 stocks were produced and titrated in VeroE6 cells as described previously (PMID 33917313). VSV-GFP31 was kindly provided by Dr. Rodriguez (CNB-CSIC). WNV-GFP recombinant virus was rescued from cloned cDNA as reported previously^30^. Trans-complemented defective reporter HCV virions (HCVtcp) were produced as described in (Steinmann et al., 2008)^52^. The hCoV-229E-GFP^53^ was kindly provided by Dr. Thiel (University of Basel) and propagated in Huh7 cells at 33oC in a controlled 5% CO_2_ environment.

### Cypopathic effect protection assays in Vero E6 and Huh7-ACE2 cells

Vero E6 or Huh7-ACE2 cell monolayers were inoculated (MOI = 0.001) in the presence of a wide range of two-fold dilutions of the crude, or partially purified extracts, or pure compounds and incubated for 72 h. Cytopathic effect and lack thereof was visualized by crystal violet staining, as previously described^33^. Untreated and solvent-treated cells were included in each plate as controls.

### Evaluation of the antiviral activity by immunofluorescence microscopy

Vero E6, A549-ACE2 or Calu3 were seeded onto 96-well plates as described above and infected in the presence of the indicated compound dose (MOI = 0.01). Twenty-four hours post infection (48 h for Calu3 cells), cells were fixed for 20 minutes at troom temperature with a 4% formaldehyde solution in PBS, washed twice with PBS and incubated with incubation buffer (3% BSA; 0.3% Triton X100 in PBS) for 1 h. A monoclonal antibody against the N protein was diluted in the incubation buffer (1:2000, v/v; Genetex HL344) and incubated with the cells for 1 h; after this time, cells were washed with PBS and subsequently incubated with a 1:500 (v/v) dilution of a goat anti-rabbit conjugated to Alexa 488 (Invitrogen-Carlsbad, CA). Nuclei were stained with DAPI (Life Technologies) as recommended by the manufacturer during the secondary antibody incubation. Cells were washed with PBS and imaged using an automated multimode reader (TECAN Spark Cyto; Austria).

All the infection experiments were performed by mixing the virus and compound dilutions 1:1 (v/v) before addition to the target cells. In the time-of-addition experiments, Vero E6 cultures were inoculated (MOI from 0.5-1) for 1 h in the presence or absence of the compounds at 37 ^o^C. Subsequently, virus-compound mixtures were left at all times, or removed and replaced with fresh 2% FBS complete media containing or not the tested compounds (see experimental scheme in Figure 8 for details). Cells were fixed 6 h post-inoculation.

### Viral RNA quantitation by RT-qPCR

A549-ACE2 cell monolayers were inoculated at MOI = 0.001 in the presence of non-toxic concentrations of the compound. Forty-eight hours later, cell supernatants were collected and heat-inactivated as described in (Smyrlaki et al., 2020)^54^, and processed directly for RT-qPCR. Alternatively, cell lysates were prepared using the Trizol reagent (Thermo Scientific) and the viral RNA content was determined by RT-qPCR using previously validated sets of primers and probes specific for the detection of the SARS-CoV-2 E gene and the cellular 18S gene, for normalization purposes. ΔCt method was used for relative quantitation of the intracellular viral RNA accumulation in compound-treated cells compared to the levels in infected cells treated with DMSO (set as 100%).

### Cytotoxicity measurement by MTT assays

Cell monolayers were seeded in 96-well plates. The day after cells were treated with a wide range of compound concentrations and forty-eight hours later they were subjected to an MTT assays using standard procedures^55^. The CC_50_ values were graphically interpolated from dose-response curves obtained with three biological replicates.

### Assessment of viral entry using retroviral pseudotypes

Retroviral particles pseudotyped with SARS-2-CoV spike envelope protein (Spp) were produced in HEK293T cells as previously described^39^ with materials kindly provided by Dr. F. L. Cosset (INSERM, Lyon) and J. M. Casasnovas and J. G. Arriaza (CNB-CSIC) for the S protein cDNA. Particles devoid of envelope glycoproteins were produced in parallel as controls.

### Statitistical Analysis

Descriptive statistics were calculated using Microsoft Excel. One-way ANOVA and *post-hoc* tests were calculated using IBM SPSS Software Package (version 26). EC_50_ and EC_90_ values were obtained employing the PROBIT regression method^56^ using IBM SPSS vs26. Synergy analysis was carried out in the web-based platform Synergy Finder (https://synergyfinder.fimm.fi/)^32^.

## Acknowledgements

This research was funded by CSIC (PIE-RD-COVID-19 ref 202040E236 to R.S., PIE-RD-COVID-19 ref. E202020E079 to P.G and U.G, and CSIC-CoV19-153 ref PIE-202080E221 to M.G-R.), EVA (European Virus Archive; grant agreement No 871029 to P.G.) and the European Union-NextGenerationEU. G.H.J-A. was supported by the Deutsche Forschungsgemeinschaft (Individual Research Grant JI 241/2-1). We thank Gema Calvo, Jennifer Moya, Laura Barbado and Enara San Sebastian for outstanding technical assistance.

## Footnotes

1 The TLC was heated (heat gun) vigorously at the extract application point before being developed; this procedure enhances Chlorophylls conversion into PheoA.

## Notes

### Competing Interest Statement

The authors have declared no competing interest.

