## Supplementary Figures for "SARS-CoV-2 fears green: the chlorophyll catabolite Pheophorbide a is a potent antiviral"

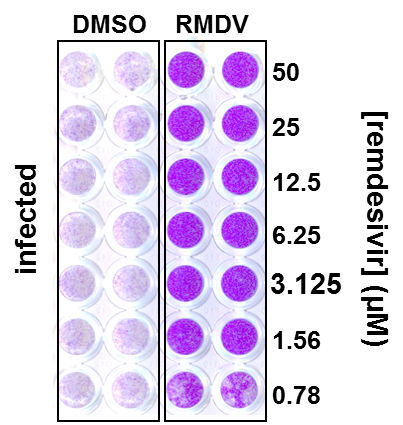

**Figure S1: Remdesivir interferes with SARS-CoV-2-induced cytopathic effect.**

Vero E6 cells were inoculated with SARS-CoV-2 at MOI = 0.001 in the presence of serial 2-fold dilutions of remdesivir (50-0.78 µM) and incubated for 72 h, time after which they were fixed and stained with a crystal violet solution. Representative image showing that interference with viral multiplication by remdesivir reveals its protective activity against virus-induced cytopathic effect at the expected concentrations.

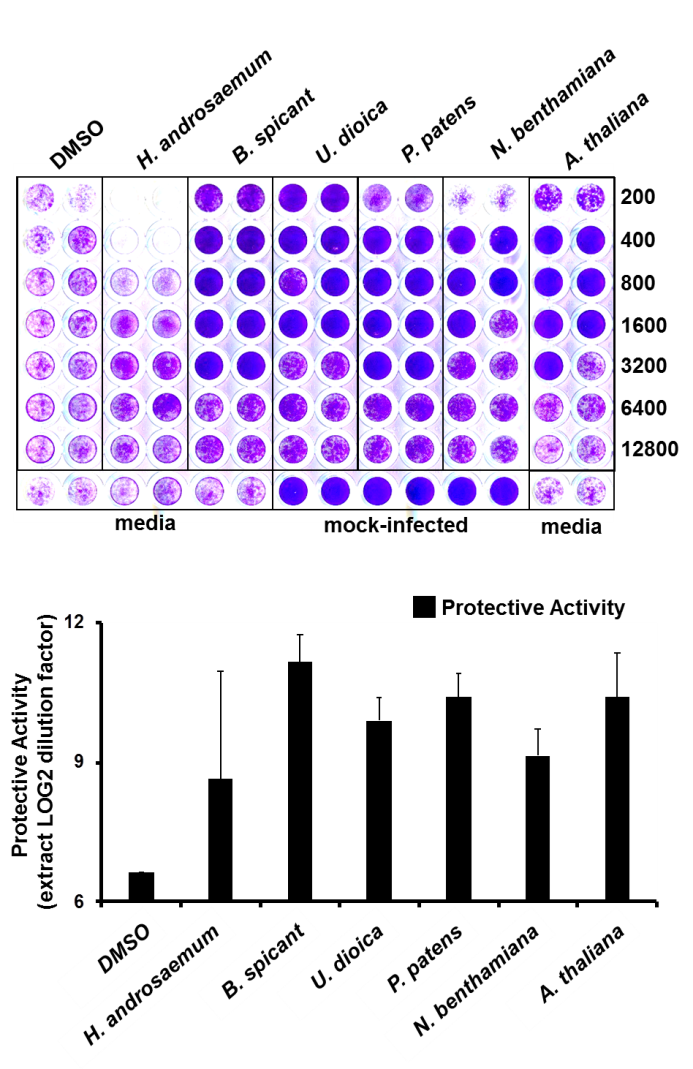

**Figure S2: Crude extracts of several plant species show antiviral potential against SARS-CoV-2 infection in cell culture.**

Vero E6 cells were inoculated with SARS-CoV-2 at MOI of 0.001 in the presence of serial 2-fold dilutions of crude extracts from sweet amber (Hypericum androsaemum), fern (Blechnum spicant), nettle (Urtica dioica), moss (Physcomitrium patens), tobacco (Nicotiana benthamiana), thale cress (Arabidopsis thaliana). Cultures were incubated for 72 h, time after which they were fixed and stained with a crystal violet solution. Mock-infected cells were used as a control of the integrity of the cell monolayer. A) Image of an experimental plate showing the dose-dependent protection of plant extracts in comparison with vehicle (DMSO)-treated cells. B) Numeric expression of the protective capacity as the log2 value of the highest dilution factor capable of full monolayer protection. Data are shown as average and mean error of two biological replicates.

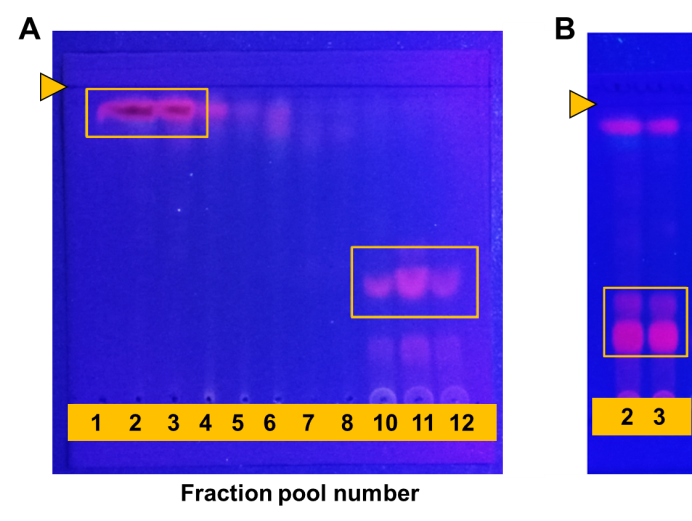

**Figure S3: TLCs of pooled fractions submitted to antiviral assays.**

A) TLC plate of chromatographic fractions under long wave ultraviolet light (365 nm), no heat. B) TLC of chlorophyll containing fractions (2 and 3) after heating the plate before being developed. Red fluorescent spots, characteristic of plant chlorophylls are squared. Solvent system AcOEt:MeOH (9:1, v/v). Arrow heads depict the solvent front.

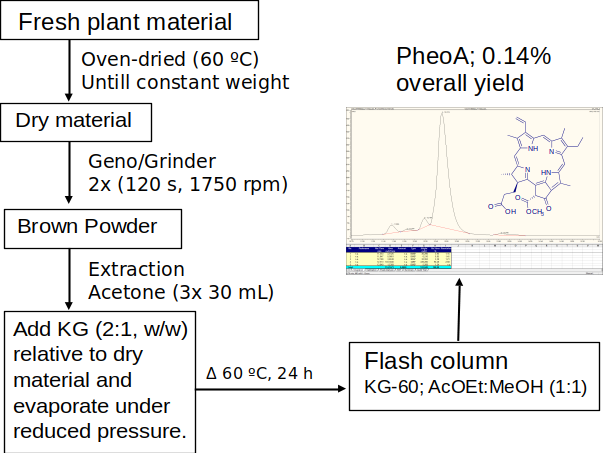

**Figure S4:** Scheme of Pheophorbide a preparation from plant material. KG, Silica gel 60.

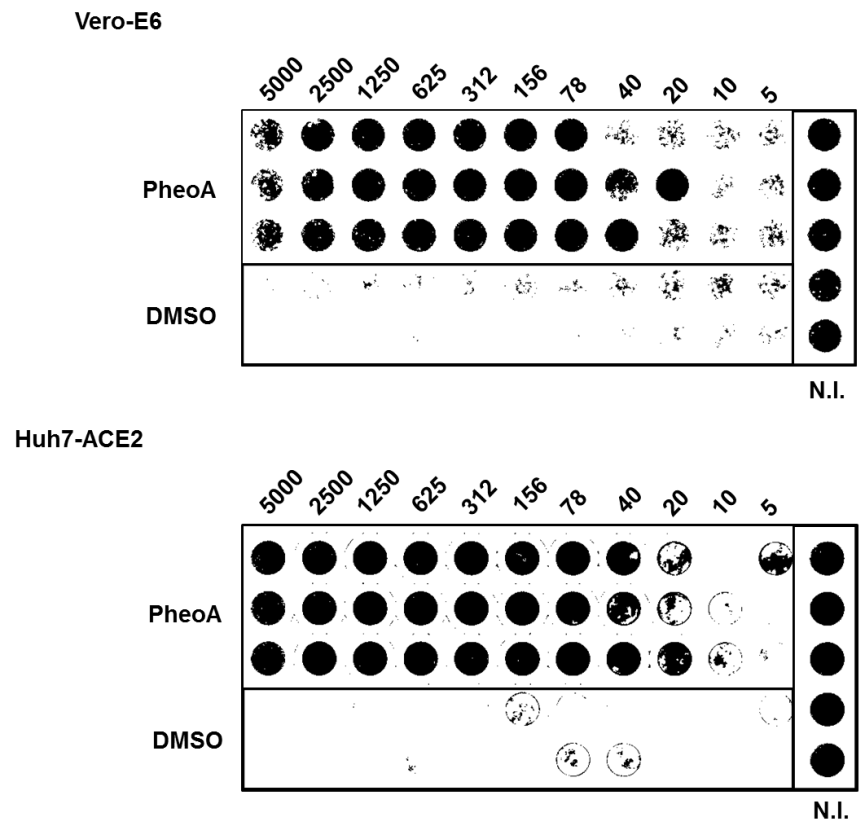

**Figure S5: Pheophorbide a protects Vero-E6 and human hepatoma Huh7 cells from virus-induced cytopathic effect.**

Vero E6 or Huh7 cells were inoculated with SARS-CoV-2 at MOI = 0.001 in the presence of the indicated PheoA concentrations. Cultures were incubated for 72 h, time after which they were fixed and stained with a crystal violet solution. Mock-infected (N.I.) cells were used as a control of the integrity of the cell monolayer.

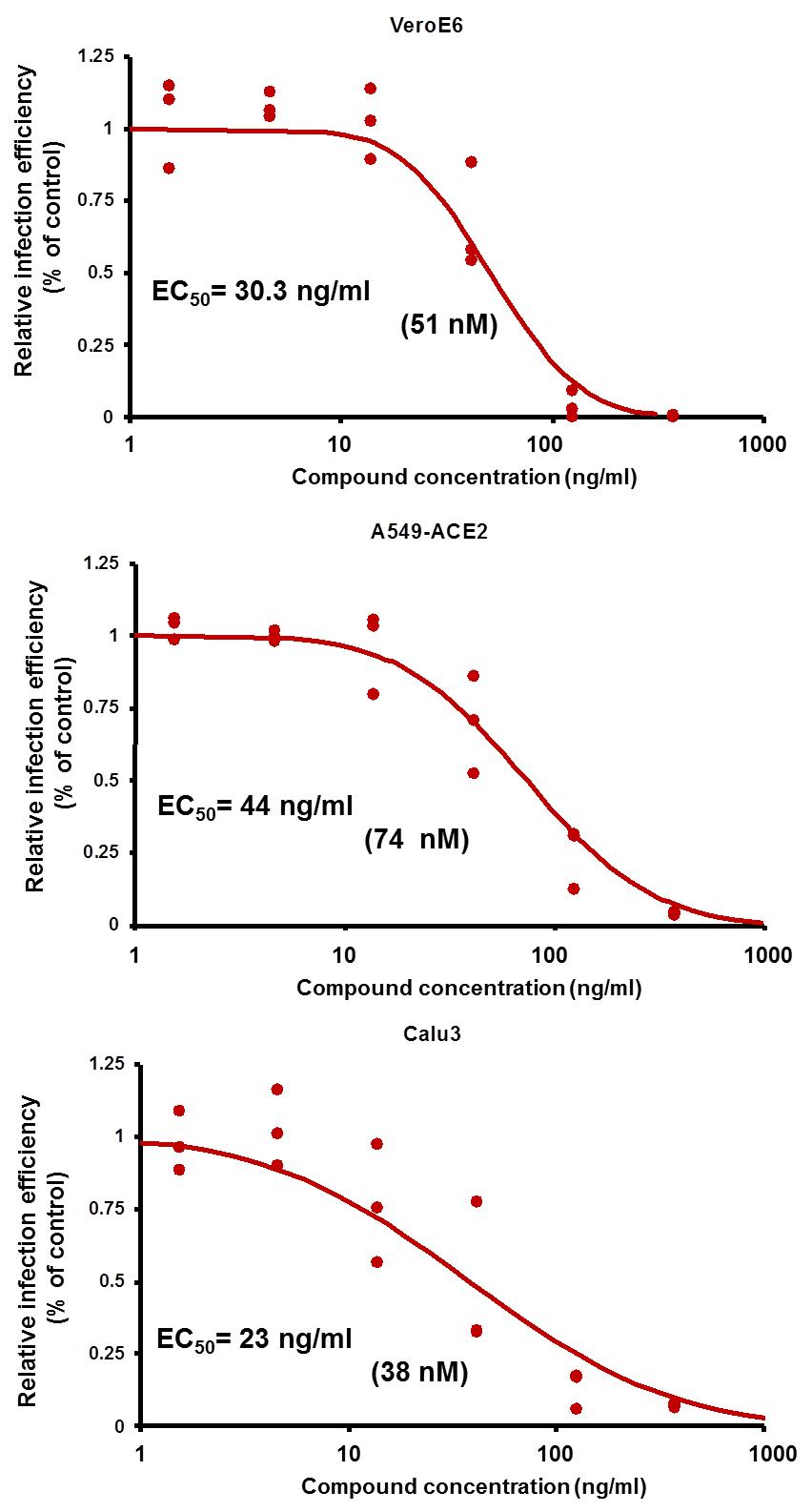

**Figure S6: Semi-synthetic PheoA preparations display antiviral activity against SARS-CoV-2.**

Semi-synthetic PheoA was produced as described in Figure S4, serially diluted and mixed 1:1 with SARS-CoV-2 preparations to achieve the indicated compound concentrations and a final multiplicity of infection of 0.005 for Vero E6 and Calu3, and 0.01 for A549-ACE2 cells. Cultures were incubated for 48 h, fixed and processed for automated immunofluorescence microscopy analysis. Relative infection efficiency data (N=3 per dose) are shown as individual data and a PROBIT regression curve (red line) using the represented values.

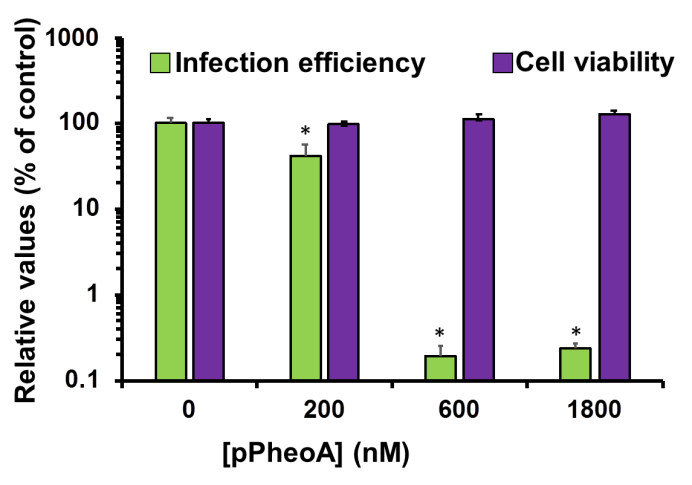

**Figure S7: Pyropheophorbide a displays antiviral activity against SARS-CoV-2.**

Commercially available pPheoA was serially diluted and mixed 1:1 with SARS-CoV-2 preparations to achieve the indicated compound concentrations and a final multiplicity of infection of 0.005 for Vero E6 and Calu3, and 0.01 for A549-ACE2 cells. Cultures were incubated for 48 h, fixed and processed for automated immunofluorescence microscopy analysis. Parallel cultures were used to determine compound cytotoxicity using an MTT assay. Relative values are expressed as the percentage of the values obtained in vehicle-treated cells and are shown as average and standard deviation (N=3 per dose). Statistical significance was estimated using one-way ANOVA and a Dunnet ́s post-hoc test (*p<0.05).

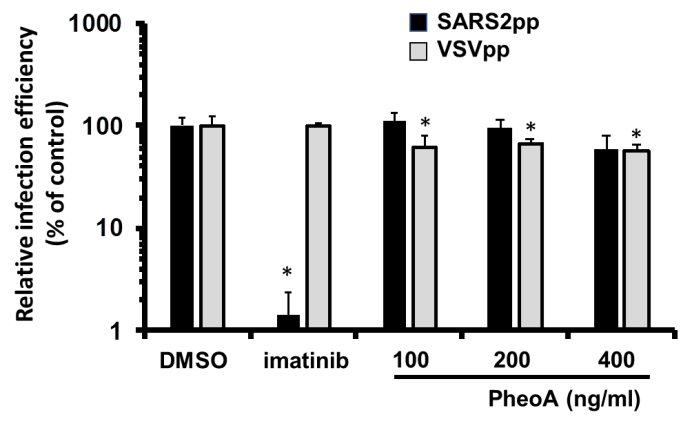

**Figure S8: SARS-CoV-2-pseudotyped retroviral vectors (Spp) are not susceptible to PheoA antiviral activity.**

Vero E6 cells were inoculated with retroviral pseudotypes bearing the SARS-CoV-2 Spike (SARS2pp) or the VSV-G glycoprotein (VSVpp) in the presence of the indicated compound concentrations. Forty-eight hours post-infection, cells were lysed and infection efficiency was estimated by the luciferase reporter gene activity. Relative infection values were calculated as percentage of the luciferase activity observed in vehicle (DMSO)-treated cells and is shown as average and standard deviation of three biological replicates (N=3). Statistical significance was estimated using one-way ANOVA and a Dunnet´s post-hoc test (*p<0.05).
